# Ontogenetic and static allometry of hind femur length in the cricket *Gryllus bimaculatus* (Orthoptera: Gryllidae) with implications for evo-devo of morphological scaling

**DOI:** 10.1101/2020.03.01.972141

**Authors:** Jakke Sameli Neiro

## Abstract

The evolution of morphological allometry or scaling is a long-standing enigma in biology. Three types of allometric relationships have been defined: static, ontogenetic and evolutionary allometry. However, the theory of the interrelationship between these three types of allometry have not been tested in Orthopterans and to a lesser extent in hemimetabolous insects. Here, the ontogenetic allometry of hind femur length in the cricket *Gryllus bimaculatus* was observed to be slightly positive as compared with a negative allometric relationship for Orthopterans in general, while the instar-specific static allometries were highly variable. The findings give support for the size-grain hypothesis in Orthoptera and indicate that ontogenetic allometries may not predict evolutionary allometries. The current model for the developmental basis of allometry derived from holometabolous insects is extended into a phylogenetic context and the potential of *G. bimaculatus* and other Orthopterans for further experiments of evo-devo of morphological scaling is discussed.

## 1. Introduction

Evolutionary developmental biology aims for a mechanistic understanding of how phenotypic diversity originates and changes throughout the course of evolution. To date, decades of research have mostly succeeded to elucidate the general mechanisms for the macroevolution of body plans and patterns, but how body shapes scale and evolve in relation to size, i.e. the evolution of morphological allometry, has remained elusive (Stern and Emlen 1999, Pélabon et al. 2014, Mirth et al. 2016, Nijhout and McKenna 2019). Nonetheless, the past decade has witnessed significant strides in elevating the study of allometry from a descriptive science of observed relative growth patterns in diverse taxa into a unified field studying the functional mechanisms that generate allometric relationships and how these respond and influence evolution (Nijhout and McKenna 2019).

### 1.1 Theory of allometry

Today, the mathematical theory of allometry is based on two separate definitions: the Huxley-Jolicoeur school (classical or narrow-sense allometry) defines allometry based on relative growth differences of traits, whereas the Gould-Mosimann school (broad-sense allometry) defines allometry as the covariation of shape and size that are regarded as separable quantities (Pélabon et al. 2014, Klingenberg 2016). In the classical bivariate setting *sensu* Huxley, the allometric parameters are estimated by linear regression on the log transformed trait values, *log(y) = log(b) + a log(x)*, where *y* is the size of a trait, x is the size of another trait (usually body size), *a* is the allometric slope and *b* is a constant (Pélabon et al. 2013, 2014, Klingenberg 2016). If the slope of the line is one, the allometric relationship is termed isometric, while hypoallometry or negative allometry and hyperallometry or positive allometry imply negative and positive slopes, respectively (Pélabon et al. 2013, 2014). Correspondingly, principal component analysis (PCA) is used to derive allometries in the multivariate setting by approximating the first principal component to be equal to the allometric slope in multivariate space (Klingenberg 2016). Conversely, the Gould-Mosimann school utilizes geometric morphometrics and multivariate regression of shape on the centroid size of the shape to derive allometric relationships (Klingenberg 2016). However, the two schools of thought are logically and biologically compatible with each other, and the main differences arise in how morphology is captured for further quantitative analysis (Klingenberg 2016). The Huxley-Jolicoeur school is based on linear measurements broadly applicable to all taxa, while geometric morphometrics is based on landmarks from rigid structures such as endo- and exoskeletons. Here, allometry is examined within the classical Huxley-Jolicoeur paradigm, since it is applicable to body structures freely movable in relation to the body (such as the allometry of appendages in relation to body size), the mathematical interrelationship of allometric relationships in different evolutionary contexts is better understood (Pélabon et al. 2013, 2014), and the current theory of the developmental basis of allometry is based on this framework (Shingleton et al. 2007, Shingleton et al. 2008, Mirth et al. 2016, Frankino et al. 2019).

### 1.2 Evolution of allometry

The evolution and evolvability of allometry has been a subject of debate ever since the concept of allometry was first defined (Gayon 2000, Frankino et al. 2005, Pélabon et al. 2014, Voje et al. 2014). Allometric relationships can be observed at different evolutionary levels, and three types have traditionally been conceptualized: static, ontogenetic and evolutionary allometry (Cock 1966, Gould 1966, Cheverud 1982, Pélabon et al. 2013, 2014). Static allometry is measured among individuals from the same population at the same developmental stage, ontogenetic allometry is measured among individuals from the same population but from different developmental stages, and evolutionary allometry is measured between populations, species or higher taxonomic levels (Fig. 1) (Cock 1966, Gould 1966, Cheverud 1982, Pélabon et al. 2013, 2014). All the three types of allometries have been observed to be relatively invariable between closely related species, which has been explained by two main hypotheses: allometric relationships represent either developmental constraints that direct evolution in morphospace, or they are the result of strong selection guaranteeing functional optimization (Gould 1966, Cheverud 1982, Frankino et al. 2005, Pélabon et al. 2014, Voje et al. 2014). The evolvability of a trait can be inferred by quantifying its variance or its actual response to selection, both of which have been recently applied to the study of allometry. Voje et al. (2014) systematically meta-analyzed more than 300 empirical estimates of static allometries and found the phenotypic variance to be low, while others have estimated the heritability to be low (Atchley and Rutledge 1980, Pavlicev et al. 2011). In contrast, experiments in the horned beetle *Onthophagus acuminatus*, the buttefly *Bicyclus anynana*, and the fruit fly *Drosophila melanogaster* have showed that ontogenetic relationships are evolvable under artificial selection, suggesting that selection is the main source of allometric invariability (Emlen 1996, Frankino et al. 2005, Egset et al. 2012, Shingleton and Frankino 2013, Bolstad et al. 2015, Stilwell et al. 2016, Houle et al. 2019). However, it is unlikely that selection on a single trait alone would maintain allometries for millions of years, and Houle et al. (2019) have noted that selection on pleitropic effects is likely the main contributing factor.

**Fig. 1.**
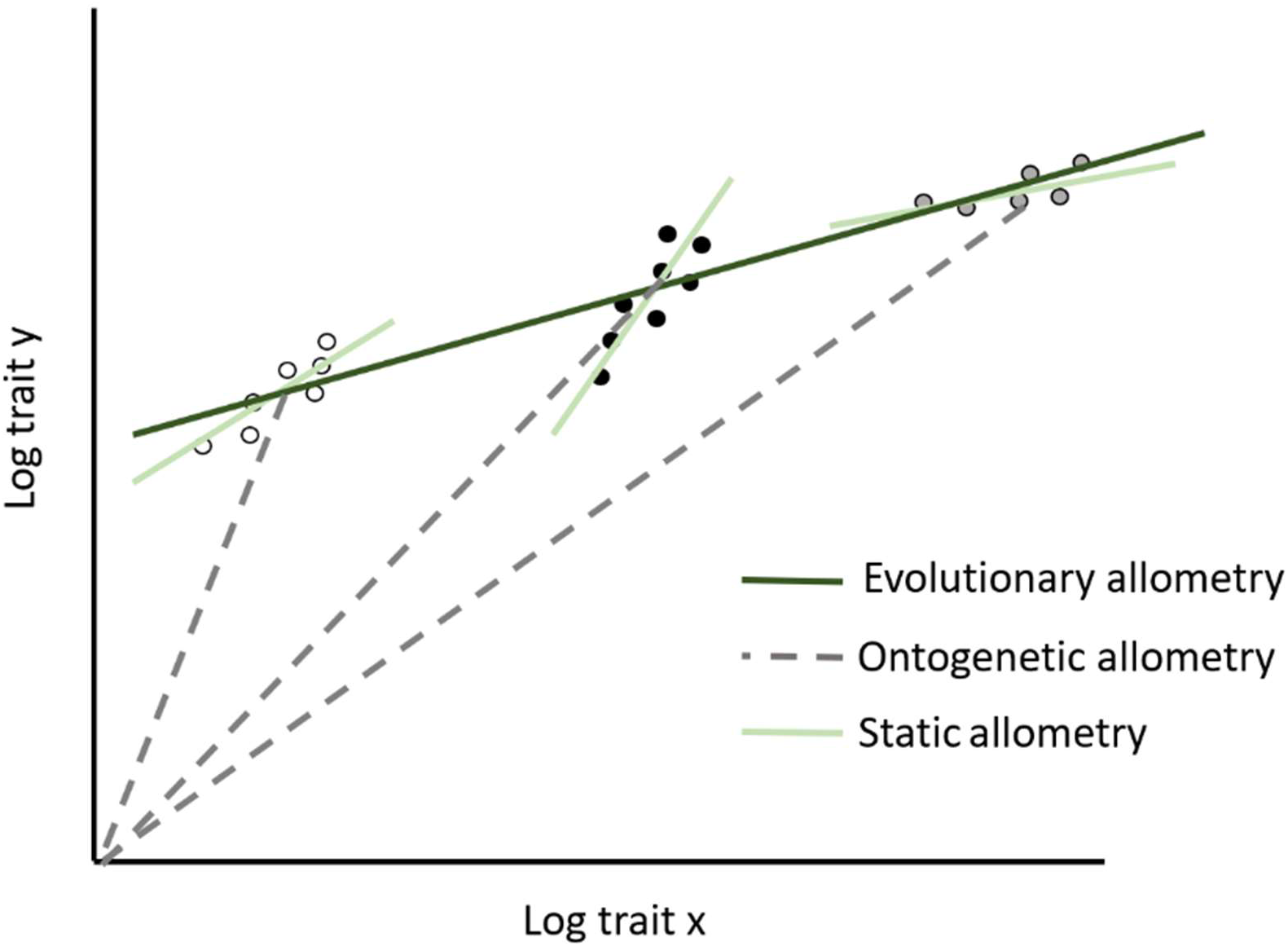
Static, ontogenetic and evolutionary allometry. Ontogenetic allometry refers to the relationship of the size of a trait *y* to the size of another trait *x* during development. Here, three different species with different ontogenetic allometries are shown. Static allometry refers to the relationship within a population of individuals from the same developmental stage in the same species, while evolutionary allometry is the relationship observed between species.

Surprisingly, the interrelationship between static, ontogenetic and evolutionary allometries have received less attention, although it has been generally acknowledged that changes in ontogenetic allometry must underlie changes in static allometry, which in turn causes transitions in evolutionary allometries, leading to the idea the three types of allometry might be used to predict each other (Gould 1966, Cheverud 1982, Pélabon et al. 2013, 2014). However, the few empirical studies on the subject have shown that the different types of allometries are not necessarily related (Cheverud 1982, Leamy and Bradley 1982, Klingenberg and Zimmermann 1992, Voje et al. 2014). Pélabon et al. (2013) were the first to formalize mathematically the interrelationship between static, ontogenetic and evolutionary allometry within the Huxley-Jolicoeur paradigm and showed that the static allometry differs from the mean ontogenetic allometry of a population if the parameters of the ontogenetic allometry and body size are correlated across individuals. Correspondingly, the evolutionary allometric slope deviates from the mean static allometric slope of selected species if the parameters of the static allometry correlate with body size across these species, indicating that the allometries would be evolvable given rather non-restrictive premises in terms of developmental biology (Pélabon et al. 2013, 2014). In addition, the mathematical theory of population-level allometries has recently been decomposed to individuals by introducing the concept of individual-level cryptic scaling relationships (Dreyer et al. 2016, Frankino et al. 2019). However, how the different types of allometry relate to each other within different taxa, the mathematical theory within the Gould-Mosimann paradigm, and the connection of population-level theory to mechanistic experiments of organ size regulation in individuals remains to be clarified.

### 1.3 Allometry and development

Interestingly, the current model for the developmental mechanism for morphological allometry is almost exclusively based on insights from holometabolous insects, especially *D. melanogaster* and horned beetles (Stern and Emlen 1999, Mirth et al. 2016, Casasa et al. 2019, Nijhout and McKenna 2019). The model posits that differential nutrition-sensitivity through the insulin/insulin-like growth factor signaling (IIS) pathway in differential organs explains differences in allometric relationships (Mirth et al. 2016, Casasa et al. 2019, Nijhout and McKenna 2019). Upon feeding, the insulin producing cells (IPCs) in the brain secrete insulin-like peptides (ILPs) that bind to insulin-like receptors (InR), which activates the phosphokinase signal transduction cascade and induce cell division and growth (Mirth et al. 2016, Casasa et al. 2017, Casasa et al. 2019). Modulation of the activity of the components in the IIS pathway, including the transcription factor Foxo and InRs, result in changes of the nutrition-sensitivity of organs and hence their relative sizes in the adult (Tang et al. 2011, Emlen et al. 2012, Snell-Rood et al. 2012, Casasa and Moczek 2018). Importantly, the IIS pathway has been observed to be involved in organ size regulation in two hemimetabolous species, the hemipterans *Nilaparvata lugens* and *Jadera haematoloma*, although the knockdown of Foxo results in opposite phenotypes (Xu et al. 2015, Fawcett et al. 2018). Overall, these findings suggest that the current mechanistic model for allometry is generally applicable within insects, but further investigation into other and more basal insect taxa would verify this hypothesis.

### 1.4 Allometry and Orthoptera

Thanks to their versatile jumping, walking, flying and chirping behavior, several Orthopteran species are classical models in functional morphology and biomechanics, including descriptive allometry (Huber et al. 1989, Fielding and DeFoliart 2008, Moradian and Walker 2008, Whitman 2008, Schöneich and Hedwig 2017, Bidau and Martínez 2018). Therefore, the biomechanics of jumping has been intensively studied, and it has been shown that based on biomechanical realities, natural selection would favor relatively longer legs in smaller species to compensate for the diminished jumping performance (Burrows 2010, Sutton and Burrows 2008). However, the opposing size-grain hypothesis holds that legs in smaller insects would be relatively shorter, as the smaller insects perceive a more rugose world in which longer legs would be a disadvantage (Levin 1992). So far, both supporting and falsifying evidence has been obtained on the matter in various insect species (Espadaler and Gómez 2001; Farji-Brener et al. 2004; Kaspari and Weiser 1999; Parr et al. 2003; Schöning et al. 2005). Recently, Bidau and Martínez (2018) found the evolutionary allometry of hind femur length with respect to body length to be negative for about 1500 Orthopteran species, thus supporting the biomechanical approach to leg length in orthopterans. However, they also noted that if the biomechanical hypothesis were to be the main functional constraint, then the static and ontogenetic allometries would also be negative in any individual orthopteran species.

In recent years, several Orthopteran species have entered the genomic and molecular arena thanks to high-throughput sequencing and functional knockdowns by RNA interference (RNAi). The two-spotted cricket *Gryllus bimaculatus* is the best known Orthopteran species from a molecular perspective thanks to two published transcriptomes (Bando et al. 2013, Zeng et al. 2013), effective RNAi both in embryos and nymphs (Mito and Noji 2008), transgenic lines and genome editing (Nakamura et al. 2010, Watanabe et al. 2012, 2017). This molecular toolkit has also been utilized to study the IIS pathway (Dabour et al. 2011), thus raising the possibility to elevate *G. bimaculatus* as a complementary model in understanding the evolutionary developmental biology of allometry. Moreover, RNAi has been successful in the Ensiferan *Acheta domesticus* and the Caeliferans *Locusta migratoria, Schistocerca americana* and *Schistocerca gregaria* (Dong and Friedrich 2005, Turchyn et al. 2011, Luo et al. 2012, Wynant et al. 2012, Santos et al. 2014), thus potentially allowing for mechanistic experiments in a wider phylogenetic context.

Taken together, here the Orthopterans are promoted as an integrative model system for the study allometry by considering the potential offered by aspects of functional morphology, evolutionary theory and molecular biology. In order to do so, the interrelationship of static, ontogenetic and evolutionary allometry in Orthoptera is clarified by estimating the ontogenetic and nymph-specific allometries of hind femur length in the foremost model species *G. bimaculatus* and comparing these to the corresponding evolutionary allometric relationship determined by Bidau and Martinez (2018). Considering these results, the validity of the biomechanical and the size-grain hypotheses are assessed as sources of selection for functional morphology. Finally, the implications of the observed allometric relationships for IIS signaling are outlined with testable mechanistic hypotheses.

## 2. Materials and methods

The laboratory stocks of *G. bimaculatus* were purchased from ReptileManiacs (ReptileManiacs Ay, Y-tunnus: 2440893-2, Tampere, Finland). The nymphal stages were maintained in medium-sized plastic boxes with a perforated lid (28 cm x 18 cm x 12.5 cm), while the adults were held in bigger terraria (37 cm x 22 cm x 18 cm). Crumpled paper was provided to reduce the risk of cannibalism. The crickets were fed JBL NovoBel Flakes, and water was provided every second day by moistening tissue paper (Katrin Classic C-Fold 2 Yellow).

The eggs were collected from the adult terrarium by collecting the tissue paper and incubating it in a small plastic box (13.5 cm x 10 cm x 7 cm) until hatching. The tissue paper bearing the eggs was moistened each day to keep the eggs hydrated. The crickets and the eggs were maintained at 30°C under 12L-12D light conditions.

The life cycle of crickets was followed for 28 days, during which 7-10 crickets were sampled each day and imaged under a stereomicroscope (Leica EZ4W) when being anesthetized with carbon dioxide. The total number of imaged crickets was 286. The body length, head width and hind femur length were measured with ImageJ. Body length is defined as the length measured from the distal end of frons in the head to the distal tip of the epiproct in the abdomen, head width as the length from the distal tip of the left eye to the right eye when viewed from the dorsal side, and hind femur length as the length from the trochanteric to the tibial tip of the third leg as by Bidau and Martinez (2018). The ontogenetic and static allometries were obtained from bivariate log-log data by linear regression.

## 3. Results

The ontogenetic allometry of hind femur length with respect to body length displayed a slight but significant hyperallometric relation (Fig. 2, Table 1). The linear model showed a good fit, indicating that the ontogenetic allometry could be well captured by a classical linear relationship in the bivariate log-log setting (Fig 2., Table 1). Interestingly, the static allometries for each instar were hypoallometric, but the variability was very high in certain instars, including even negative values, and the linear model had generally a poor fit (Fig. 2, Tab. 1).

**Fig. 2.**
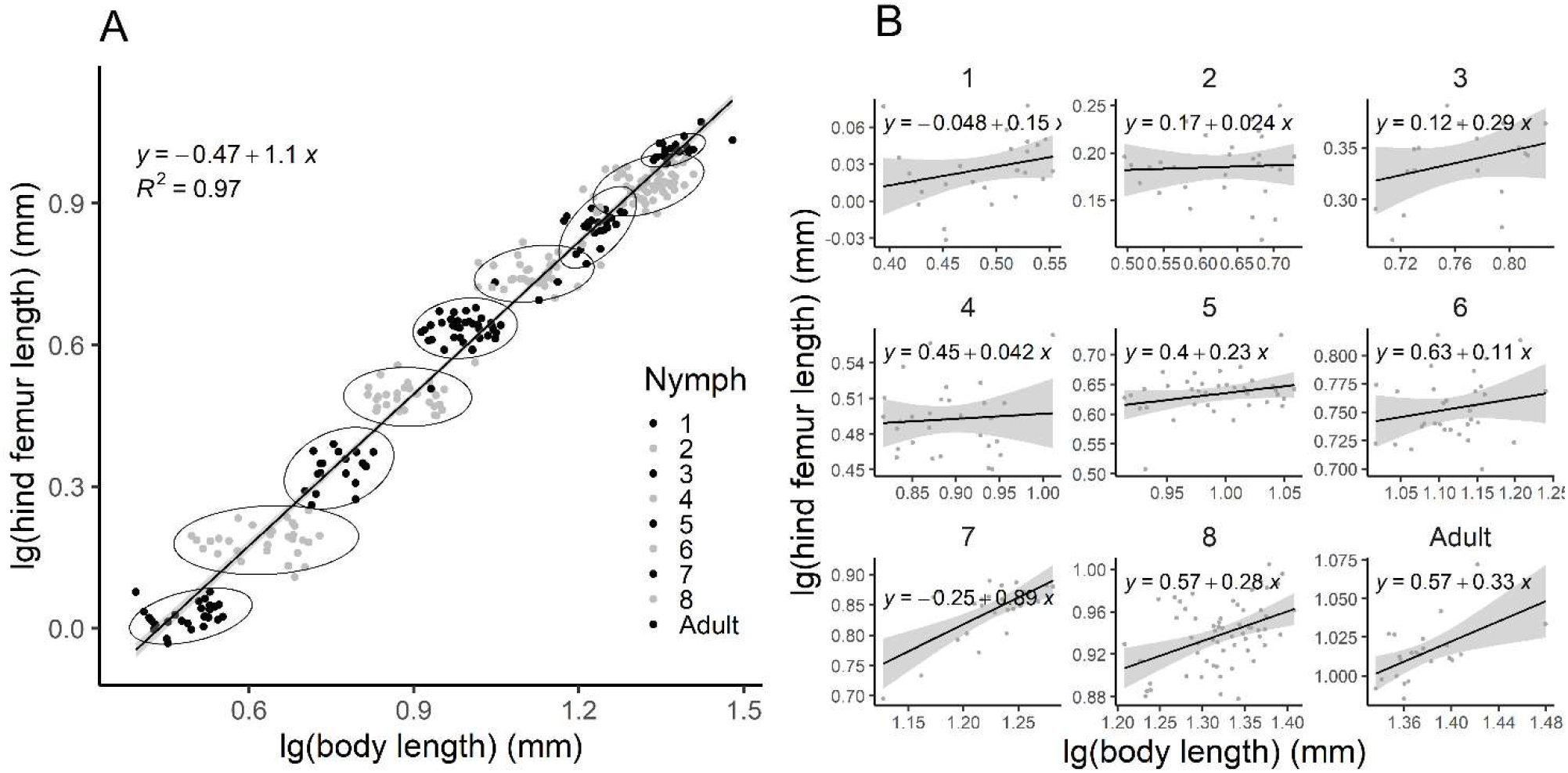
A. Ontogenetic allometry of hind femur length with respect to body length. B. Instar-specific static allometries of hind femur length with respect to body length.

**Table 1.** Ontogenetic and static allometry of hind femur length (y) with respect to body length (x) (log_10_(y) = alog_10_(x) + log_10_b). Model refers to the type of allometry measured: ontogenetic or stage-specific static allometry. *n* is the number of measured individuals, *R^2^* is the coefficient of determination, *a* is the allometric slope and *log_10_b* is the allometric constant. Confidence intervals (CI 95%) are given in parentheses.

| Model | n | $R^2$ | a | $\log_{10}b$ |
| --- | --- | --- | --- | --- |
| Ontogenetic | 286 | 0.97 | 1.07 (1.05, 1.09) | -0.47 (-0.49, -0.44) |
| Static 1<sup>st</sup> stage | 27 | 0.04 | 0.15 (-0.06, 0.37) | -0.04 (-0.16, 0.06) |
| Static 2<sup>nd</sup> stage | 29 | 0.04 | 0.02 (-0.16, 0.21) | 0.16 (0.05, 0.28) |
| Static 3<sup>rd</sup> stage | 19 | 0.04 | 0.28 (-0.17, 0.74) | 0.12 (-0.23, 0.46) |
| Static 4<sup>th</sup> stage | 29 | 0.03 | 0.04 (-0.18, 0.27) | 0.45 (0.25, 0.66) |
| Static 5<sup>th</sup> stage | 37 | 0.04 | 0.23 (-0.05, 0.50) | 0.40 (0.13, 0.68) |
| Static 6<sup>th</sup> stage | 32 | 0.008 | 0.11 (-0.09, 0.31) | 0.63 (0.41, 0.86) |
| Static 7<sup>th</sup> stage | 27 | 0.44 | 0.89 (0.49, 1.28) | -0.25 (-0.73, 0.23) |
| Static 8<sup>th</sup> stage | 60 | 0.17 | 0.28 (0.12, 0.43) | 0.57 (0.36, 0.77) |
| Static adult | 23 | 0.28 | 0.32 (0.11, 0.54) | 0.57 (0.26, 0.87) |

Correspondingly, the ontogenetic allometry of hind femur length with respect to a sclerotized structure, head width, was clearly positive and close in value to the allometry with respect to body length (Fig. 3, Table 2). Similarly, the static relationships were hypoallometric except for the 7th stage, and variability was high as when measured with respect to body length (Fig. 3, Table 2). Nevertheless, the linear model had a better fit than when measured with respect to body length, and at least the static relationship in the 8th stage and adults can be considered to be significantly negative (Fig. 3, Table 2).

**Fig. 3.**
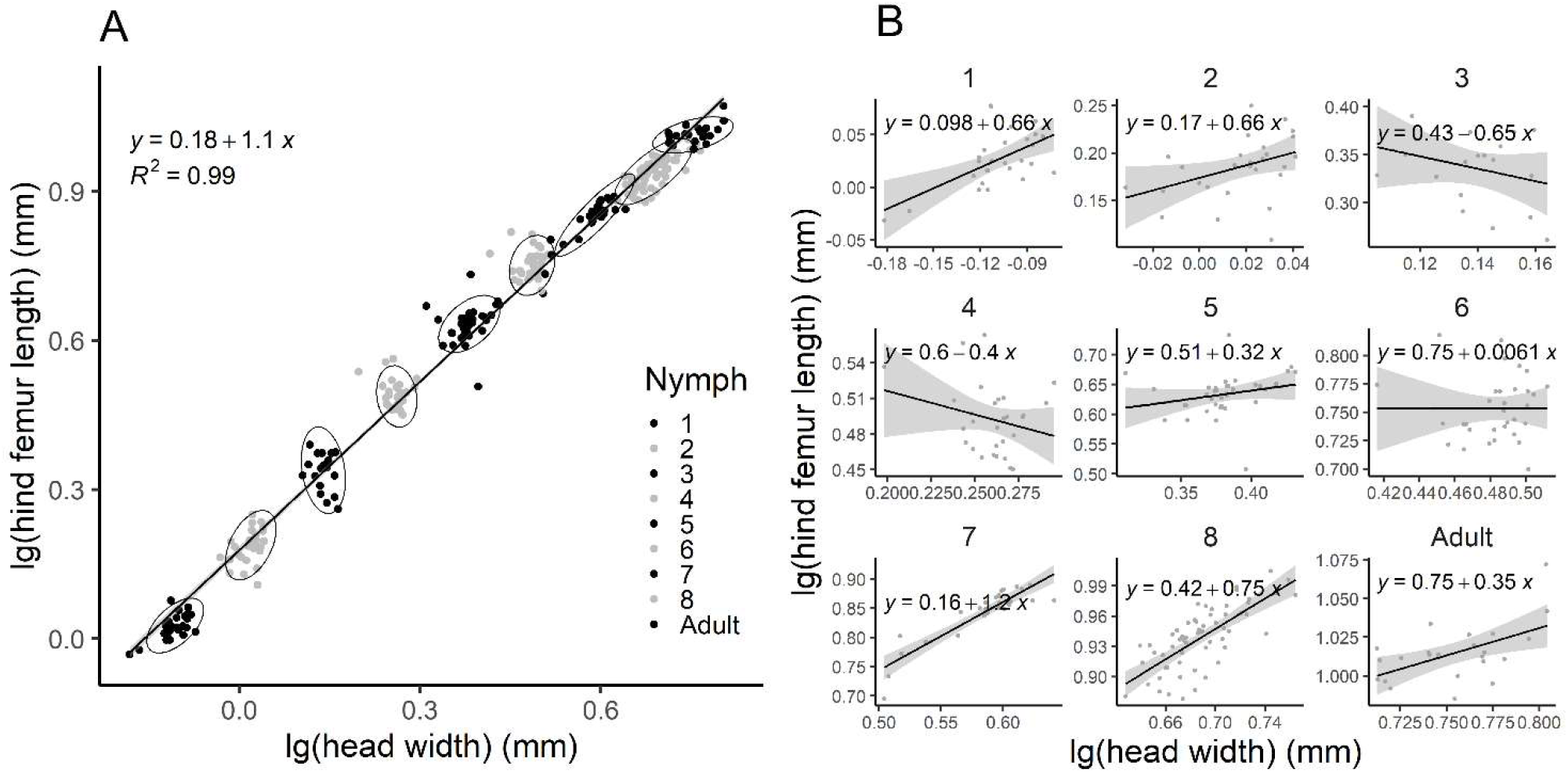
A. Ontogenetic allometry of hind femur length with respect to head width. B. Instar-specific static allometries of hind femur length with respect to body length.

**Table 2.** Ontogenetic and static allometry of hind femur length (y) with respect to head width (x) (log_10_(y) = alog_10_(x) + log_10_b). Model refers to the type of allometry measured: ontogenetic or stage-specific static allometry. *n* is the number of measured individuals, *R^2^* is the coefficient of determination, *a* is the allometric slope and *log_10_b* is the allometric constant. Confidence intervals (CI 95%) are given in parentheses.

| Model | n | $R^2$ | a | $\log_{10}b$ |
| --- | --- | --- | --- | --- |
| Ontogenetic | 286 | 0.99 | 1.13 (1.11, 1.15) | 0.18 (0.17, 0.18) |
| Static 1<sup>st</sup> stage | 27 | 0.32 | 0.66 (0.29, 1.03) | 0.10 (0.06, 0.14) |
| Static 2<sup>nd</sup> stage | 29 | 0.12 | 0.66 (0.04, 1.29) | 0.17 (0.16, 0.19) |
| Static 3<sup>rd</sup> stage | 19 | 0.25 | -0.65 (-1.78, 0.49) | 0.43 (0.27, 0.59) |
| Static 4<sup>th</sup> stage | 29 | 0.03 | -0.39 (-1.02, 0.22) | 0.60 (0.43, 0.76) |
| Static 5<sup>th</sup> stage | 37 | 0.03 | 0.32 (-0.13, 0.77) | 0.51 (0.34, 0.68) |
| Static 6<sup>th</sup> stage | 32 | 0.03 | 0.01 (-0.52, 0.53) | 0.75 (0.49, 1.01) |
| Static 7<sup>th</sup> stage | 27 | 0.81 | 1.17 (0.94, 1.40) | 0.16 (0.03, 0.30) |
| Static 8<sup>th</sup> stage | 60 | 0.53 | 0.75 (0.57, 0.93) | 0.42 (0.29, 0.55) |
| Static adult | 23 | 0.27 | 0.35 (0.11, 0.60) | 0.75 (0.56, 0.93) |

## 4. Discussion

### 4.1 Ontogenetic allometry differs from evolutionary allometry, static allometries are inconclusive

In this article, the interrelationship of static, ontogenetic and evolutionary allometry of the third hind femur was investigated for the first time in Orthoptera. Interestingly, the ontogenetic allometry of hind femur length with respect to both body length and head width in the cricket *G. bimaculatus* was clearly linear and isometric or slightly positive and not negative as reported for the evolutionary allometry across Orthopterans in general or Gryllidae in particular (Bidau and Martínez 2018). The result follows the only other similar comparison in hemimetabolous insects by Klingenberg and Zimmermann (1992), who noted that the evolutionary allometry in the water strider genera *Gerris* and *Aquaticus* was markedly different from both static and ontogenetic allometries. Also, Voje et al. (2014) noted a similar trend overall based on data from both holometabolous insects and vertebrates, and thus the findings for Orthoptera are consistent with previously reported empirical patterns.

However, the static allometries were highly variable for almost all nymphal instars, which in part is explained by the lower instar-specific sample size than for overall ontogenetic allometry, but all the same the static relationships are more variable than expected from earlier studies. The abdomen of crickets is known to be highly telescopic, which apparently confounds the static relationships with respect to body size in individual instars, as the observed variance for body size is far greater than for hind femur width. The static allometries with respect to head width are slightly less variable, but still no clear relationship can be distinguished for most instars. The high variability of static allometries could be attributable to the experimental setup and inexact measurement of body dimensions from microscopic images, but a few observed patterns are inconsistent with this as the sole explanation. Firstly, measurement error would be equally distributed to all measured linear dimensions in all instars, but the results show that the static allometries follow the linear model better in later instars. As the linear measurements were captured from images taken with different magnifications, the size of different body structures was equal in proportion to overall image size in all instars. Secondly, the 7^th^ instar exhibits the least variance and the best fit to a linear static allometry when measured with respect to either body size or head width. Thirdly, when hind femur length is measured with respect to head with, the best linear allometries can be seen in the first and last instars, while the middle instars show the poorest fit to the linear model. Therefore, it is concluded that the observed pattern for static allometries arise mostly from biological factors and not from experimental bias.

The mathematical model of linear allometry by Pélabon et al. (2013) predicts that in order to observe allometric relationships, the range of values for both compared traits needs to be large enough to overcome individual variation. Here, it seems that the instars of *G. bimaculatus* do not follow this mathematical requirement, and therefore static allometries measured for individual instars are inconclusive. Klingenberg and Zimmermann (1992) measured the static allometries for five instars in waterstriders, and it might be that in crickets with eight instars a too small proportion of the overall trait size distribution is discretized into a single instar. Nonetheless, the cricket *G. bimaculatus* undergoes the largest morphological internymphal transition between the sixth and seventh instar, a transition that also has been noted to have a high mortality in other cricket species (Zhemchuzhnikov and Knyazev 2011). Therefore, the interestingly neat static relationship observed for the seventh instar could reflect increased selection during this metamorphotically critical period in the development of *G. bimaculatus*. However, further investigation of static allometries in different hemimetabolous insects and arthropods would shed more light on these potential peculiarities of measuring allometries in molting organisms. Furthermore, although the static allometries are highly variable, most of the observed relationships are significantly hypoallometric, suggesting that the static allometries may differ from the ontogenetic allometry and are in fact closer to the negative evolutionary allometry observed for Grylliade and Orthopterans in general (Bidau and Martínez 2018).

### 4.2 Ontogenetic allometry reflects different selection regimes

The clear slightly positive ontogenetic allometry in *G. bimaculatus* could suggest a trade-off between the size-grain hypothesis predicting a positive allometry of hind femur length and the biomechanical optimization of jumping performance predicting a negative allometry of hind femur length in Orthoptera (Levin 1992, Sutton and Burrows 2008, Burrows 2010, Bidau and Martínez 2018). This could indicate that the relative fitness advantage of jumping performance in the laboratory stock of *G. bimaculatus* is smaller than that for Gryllidae and Orthoptera in general. In contrast, the relative fitness advantage of walking would be higher, which is not implausible given that crickets in captivity rarely have the space to jump but walk continuously. However, the size-grain hypothesis would predict that the static allometries would also be isometric or positive, but no such pattern is evident, possibly due to caveats discussed above.

The mathematical model by Pélabon et al. (2013) predicts that the evolutionary allometry would be more negative than the static or ontogenetic allometries if the allometric slope correlated negatively with body size in individuals, which would explain the observed positive ontogenetic allometry and negative static and evolutionary allometries. In other words, this would mean that crickets with a larger final body size would also have a more negative individual ontogenetic slope, which could be theorized to be the result of differences in locomotory behavior in small and large crickets. Crickets with a genetic predisposition of becoming larger may jump less and walk more as adults, and thus the size-grain selection pressure would be stronger in larger individuals and the allometric slope would correlate negatively with body size. However, this would also indicate that larger species would also have more negative ontogenetic allometries, and therefore further investigation into the ontogenetic allometries of different Orthopteran species would further test the predictions of the current model by Pélabon et al. (2013). Several Orthopteran species have been successfully cultivated, and therefore the order possesses a good potential for becoming a good experimental system for validating the current theory of the evolution of morphological allometry (Hinks and Erlandson 1994).

Interestingly, research in the evolution of allometry has often implicitly assumed that selection is uniform throughout development, and the interaction between natural selection and development has not been fully acknowledged (Pélabon et al. 2013, 2014, Voje et al. 2014). The selective force for a given trait may be stronger earlier in development and may therefore select individuals on certain developmental trajectories, and therefore observations of adults only may not reveal the primary source of selection. As growth in Orthopterans along with other hemimetabolous insects proceed through instars, developmental stages may easily be chosen and studied separately to assess how the selection regime changes during development to further understand the underlying cause of the observed allometric relationship.

### 4.3 Developmental model for static, ontogenetic and evolutionary allometry

The current developmental model for morphological allometry holds that differences in nutrition-sensitivity between organs accounts for differences is allometric relationships (Stern and Emlen 1999, Mirth et al. 2016, Casasa et al. 2019, Nijhout and McKenna 2019). Organs with a steeper allometric slope are more nutrition-sensitive and show a higher increase in IIS signaling upon an increase in the level of nutrition. However, the current model has only been explicitly formulated in the context of ontogenetic allometry and the implications for static and evolutionary allometry have not been outlined (Pélabon et al. 2013, 2014, Mirth et al. 2016, Casasa et al. 2019, Nijhout and McKenna 2019). If the current model is extended into a static context, it would indicate that the relative nutrition-sensitivity and nutrition-induced IIS signaling of a given organ would be higher in individuals with a more hyperallometric relationship between organ size and a reference (Fig. 4). Here, the relative nutrition-sensitivity of an individual organ would be measured as the increase in IIS signaling upon feeding as compared with the corresponding increase in a number of reference organs in the same individual. Similarly, if the model is extended to the evolutionary context, the relative nutrition-sensitivity and nutrition-induced IIS signaling would be higher in species with a more hyperallometric relationship between organ size and a reference (Fig. 4).

**Fig. 4.**
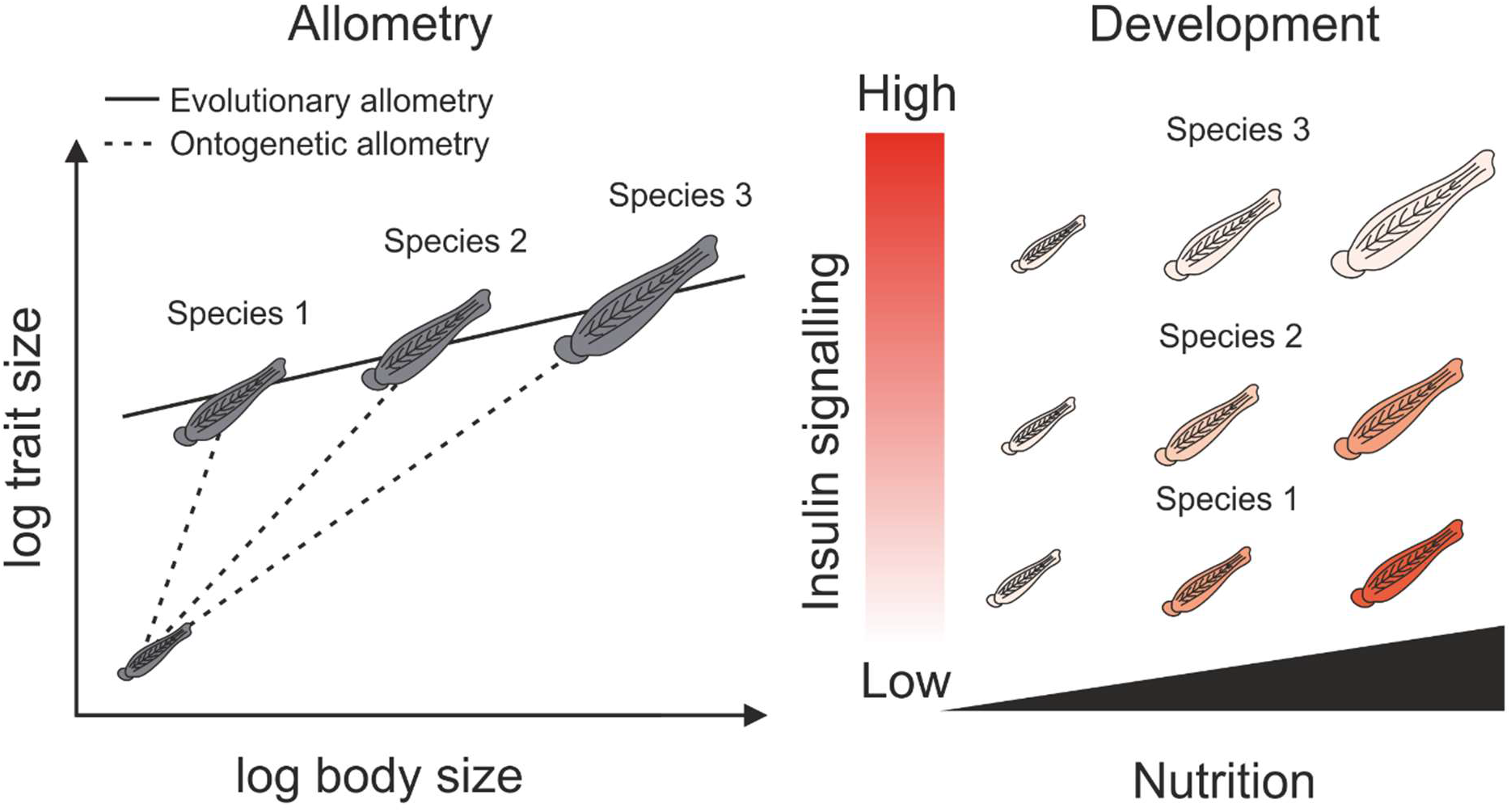
Hypothetical model of the developmental basis of static, ontogenetic and evolutionary allometry based on Mirth et al. (2016). In individuals (static allometry) or species (evolutionary allometry) with differences in the ontogenetic allometry, the individuals or species with a steeper ontogenetic slope are hypothetically more nutrition-sensitive. This would indicate that upon an increase in nutrition, the relative increase in IIS signaling would also be higher. Conversely, in individuals or species with a shallower slope, the relative nutrition-sensitivity and IIS response would be smaller.

Interestingly, Thompson (2019) extended recently the study of allometry into Orthopterans by investigating the diet-induced plasticity of static allometry of leg length in the grasshopper *Melanoplus sanguinipes*. He observed that grasshoppers fed a low-nutrient diet developed shorter legs in relation to body size compared to individuals fed a high-nutrient diet. In other words, nutritional deprivation resulted in a more negative allometric slope, which is in concordance with the prediction of the developmental model. Similar studies in other Orthopteran species would reveal whether this is a general trend within the order. Furthermore, experiments varying the amount of nutrition in different instars would hypothetically lead to a non-linear allometry being the result of linear allometries with different slopes during development. Alternatively, reduced phenotypic plasticity in later instars might reveal critical periods for forming morphological allometries during development.

For molecular studies, RNAi and RNA-sequencing (RNA-seq) in *G. bimaculatus* would reveal whether the observed positive allometry agrees with the developmental model based on data from holometabolous insects. Previously, InR, chico and Foxo have all been successfully knocked down globally with RNAi, leading to changes in body size overall, but potential changes in allometric relationships were not measured (Dabour et al. 2011). Hypothetically, global inhibition of the IIS pathway would lead to a more negative allometry for hind femur length. IIS signaling in the hind femur could be measured with RNA-seq or quantitative PCR to assess whether the signaling response upon feeding is slightly higher or equal in the hind femur as compared with head width. Furthermore, comparative studies in other Orthopteran species with a molecular toolkit (A. *domesticus, L. migratoria, S. americana* and *S. gregaria)* would give the possibility to empirically test the predictions of the developmental model extended here to an evolutionary context.

### 4.4 Conclusions

Here, a slightly positive ontogenetic allometry was observed for hind femur length in the field cricket *G. bimaculatus* as opposed to the negative evolutionary allometry measured in Orthoptera. The result supports the size-grain hypothesis and entails further testable hypotheses. The current developmental model of allometry is based on holometabolous insects, *D. melanogaster* and horned beetles, and here *G. bimaculatus* and other Orthopterans are acknowledged as a good potential experimental system to integrate the developmental model with the theory of the evolution of morphological allometry.

## Acknowledgements

The study was conducted as a side project of a Master’s thesis at the University of Helsinki, funded by Suomen Biologian Seura Vanamo ry (Pro gradu-apuraha), Suomen Hyönteistieteellinen Seura (Pro gradu - apuraha), and Entomologiska föreningen i Helsingfors - Helsingin hyönteistieteellinen yhdistys (Pro gradu - stipendium).

